# High speed functional imaging with a microfluidics-compatible open-top light-sheet microscope enabled by model predictive control of a tunable lens

**DOI:** 10.1101/2025.07.23.666439

**Authors:** W. Alexander Calhoun, Sihoon Moon, Lucinda Peng, Hang Lu

## Abstract

Functional fluorescence imaging of small, transparent organisms has proven to be a powerful tool for understanding the development and function of intact nervous systems at the cellular level, particularly when combined with microfluidic tools that enable precise manipulation of sensory cues. Light-sheet fluorescence microscopy has a number of advantages for functional imaging, including efficient axial sectioning, reduced photobleaching, and high speed compared to confocal methods, but many configurations with ideal properties for efficient functional imaging place constraints on sample access and geometry that preclude their use with microfluidics. We present an open-top light-sheet microscope that uses a simplified inverted water immersion interface coupled with tunable lens remote focusing to achieve high-speed, multichannel 3D imaging of specimens in conventional microfluidics. We use model predictive control to efficiently optimize drive signals for an electrically tunable lens, providing reliable, camera-limited scanning of the imaging volume. We then demonstrate the utility of this approach by recording calcium activity from *C. elegans* at volume rates up to 20 Hz, including both whole-brain response to chemical stimulation and compartmentalized dendritic response of the PVD neuron to mechanical stimulation. Our approaches are flexible and inexpensive to implement, and could be adapted to improve performance and sample compatibility for a wide range of imaging techniques.

## 2 Introduction

Light-sheet microscopy has proven to be a powerful tool for functional imaging of nervous systems, particularly when combined with genetically encoded fluorescent probes[1][2][3][4]. By using separate excitation and detection paths, light-sheet microscopy allows 3D imaging at high speed and with lower photobleaching than typical confocal methods. This technique is well-suited for use in small, transparent organisms, including *Drosophila* and zebrafish larvae and the nematode worm *C. elegans*, where it can provide detailed insights into intact nervous systems with single-cell resolution. However, this comes at the cost of complexity in design, and many configurations of light-sheet microscopes require specialized sample mounting or constrain sample size, which in turn limits the range of experimental preparations for which they are suitable. One of the advantages of small, transparent organisms as model systems for neuroscience is the ability to precisely manipulate their sensory environment using microfluidics [5][6]. However, these techniques are geometrically incompatible with many light-sheet configurations, frequently due to the need for objective lenses to surround a sample. Two categories of designs have emerged that address this issue. Open-top light-sheet microscopes with orthogonal objectives have been described, which specifically offer compatibility with microfluidics and other closed, planar samples [7][8][9][10], but these are usually geared toward large-specimen and high-throughput imaging. Single-objective oblique plane microscopes such as SCAPE [11][12], on the other hand, avoid lens geometry conflicts altogether by imaging and forming the light-sheet through the same lens. These designs have excellent performance for high-speed volumetric imaging, but are built around complex and tightly constrained optical designs, which may lose some of the flexibility of fully separated detection and excitation paths. Thus, there remains a need for simple and efficient open-top designs for high-speed functional imaging, which can be adapted to meet experimental needs.

Volumetric imaging with a light-sheet requires some method of sweeping the image plane through the sample (Fig. 1a). In its simplest form, this is accomplished by moving the sample itself while keeping the light-sheet and detection focus static. While this provides the most reliable image quality and largest potential field of view for an open-top design, it is also the slowest method. High speed functional imaging requires some way of moving the focal plane and light-sheet position quickly and synchronously. This is most commonly accomplished by using actuated micromirrors to scan the light-sheet and a piezoelectric mount to physically move the detection objective. Micromirror beam steering is fast and reliable, but piezo actuation of the detection objective is substantially limited by the inertia of the usually bulky detection objective. As the acquisition speeds of scientific cameras have improved, this has become a major limiter of achievable volume rates. Fast moving objective lenses are also problematic for liquid immersion. Maintaining immersion with an open-top interface requires a flexible seal that can accommodate the movement, and fast oscillation of the lens in the immersion medium risks mechanically disturbing the sample. A different approach is to dynamically modify the optical path to change the effective focal distance of the detection objective, referred to as remote focusing [13]. Remote focusing can not only be made much faster than objective lens scanning, it also simplifies an open-top design by eliminating the need for moving optics at the sample.

Remote focusing at volume rates up to 60 vol/s has been demonstrated using an electrically tunable lens (ETL)[14], which uses a liquid-filled membrane as a shape-changing optical element (Fig. 1b). This approach provides a simple, fast, and relatively inexpensive way of refocusing an imaging system, in the simplest case merely by adding the ETL near the back focal plane of the objective lens. The addition of the lens does introduce some spherical and chromatic aberration to the imaging system, but it has proven useful for low NA imaging and applications where speed is favored over resolution [15][16][17][18][19]. However, while an ETL module is capable of rapidly changing focal length, the liquid lens has complex response dynamics that make it difficult to use effectively for focus scanning at high speed. Light-sheet microscopy requires precise synchronization of the focal plane and light-sheet position to form an image due to the shallow depth of field of microscopy lenses, especially as detection NA increases. Without some form of signal preconditioning, a sudden step causes high frequency ringing and considerable overshoot which must be allowed to settle to form a focused image. This is typically solved through low-pass filtering of the drive signal. This slows the response time and risks negating the speed advantage of using an ETL for remote focusing, though a manually tuned initial overdrive has been shown to reduce the rise time. A more systematic approach is to use a model of ETL dynamics to optimize the drive signal [20].

A now common approach to optimizing control of a complex, but well-characterized, system is using model predictive control (MPC) [21]. Given a desired output as a reference, MPC produces optimum input signals to the system by minimizing the error between the desired and simulated future output of the system over a finite time horizon. This means that the choice of input at each time point is made with consideration of its affect on future behavior, which is useful when controlling systems that have complex or long-tailed responses to small, sudden changes in input. While MPC is most powerful when incorporating live feedback to ensure the simulated and real system states match, it makes an effective open-loop optimizer when no outside perturbations to the system state are expected and the model sufficiently captures true behavior. MPC can also incorporate constraints into the optimization, for example the minimum and maximum drive current for an ETL, preventing control failure or device damage when operating near these limits. Finally, MPC can anticipate future maneuvers when they are known in advance, as is the case for a predetermined scan waveform, which prevents sudden adjustments from reducing future controllability. These advantages are easily realized with readily available software since the parameterization lies mostly in providing an accurate model.

In this work, we constructed an open-top light-sheet microscope with a simple inverted sample interface and ETL remote focusing. We then used a black-box linear system identification approach to model the tunable lens response dynamics and used this empirical model to generate optimized drive signals using model predictive control. With this approach, achieved camera-limited volume rates of 10-20 Hz with sufficient resolution and sensitivity for functional imaging of *C. elegans* under microfluidic manipulation for several applications. The volume rate was determined by the camera acquisition speed coupled with the application-specific slice thickness required for sufficient spatial sampling.

## 3 Results

### 3.1 Design of an open-top light-sheet microscope with remote focusing for functional imaging with microfluidics

Soft lithography microfluidics is a powerful tool for manipulating small organisms such as *C. elegans*, allowing control of their mechanical and chemical environment[5][6]. However, it presents several challenges for imaging with light-sheet microscopy. Conventional microfluidic devices consist of a thick block of PDMS with microchannels embedded in the surface bonded to a rigid substrate, such as a coverslip. The microscope must be able to image across this barrier and into the channel containing the sample in order to allow pressurization (e.g. for pneumatic control or fluid manipulation) and chemical delivery [22][23]. This can be a problem for imaging at an oblique angle, as any refractive index mismatch will cause the substrate to act as a prism and introduce significant optical aberrations[8] (Supplementary Fig. 3). The simplest solution is to use water immersion and a material such as fluorinated ethylene propylene (FEP), commonly used in capillary tubes for mounting suspended samples. We found that, while less reliable than glass, corona treated FEP film will bond to PDMS under sufficient heat and pressure and can be used as the substrate for microfluidic devices, including devices with pressure actuated valves up to at least 60 PSI, sufficient for almost all applications. Our light-sheet sample interface is therefore designed to assume a water/FEP optical path to the sample without additional correction.

Open-top imaging with microfluidics also requires sufficient separation between the sample and objective lenses [7]. Unfortunately, most objective lens pairs suitable for functional light-sheet microscopy partially surround their shared focal point, making it inaccessible to samples that cannot be suspended between them [2]. We avoid this problem by using an asymmetric pair of objective lenses, a 10x/0.3NA (10X Nikon CFI Plan Fluorite) for illumination and a 40x/0.8NA (40X Nikon CFI APO NIR) for detection. This specific pair provides an effective working distance of 0.5 mm beyond the plane of contact, which is more than sufficient to image through the 50-150 *µ*m FEP film. A small 3D printed fitting maintains water immersion between the objectives and sample and is filled by syringe through connected tubing (Fig. 1c, Supplementary Fig. 2). This allows the sample to be supported on a standard XY stage without requiring it to be submerged in the immersion medium. The advantage of this configuration, thus, is that any arbitrary microfluidic design can be used to manipulate the sample as long as the barrier is index-matched to water.

**Figure 1:**
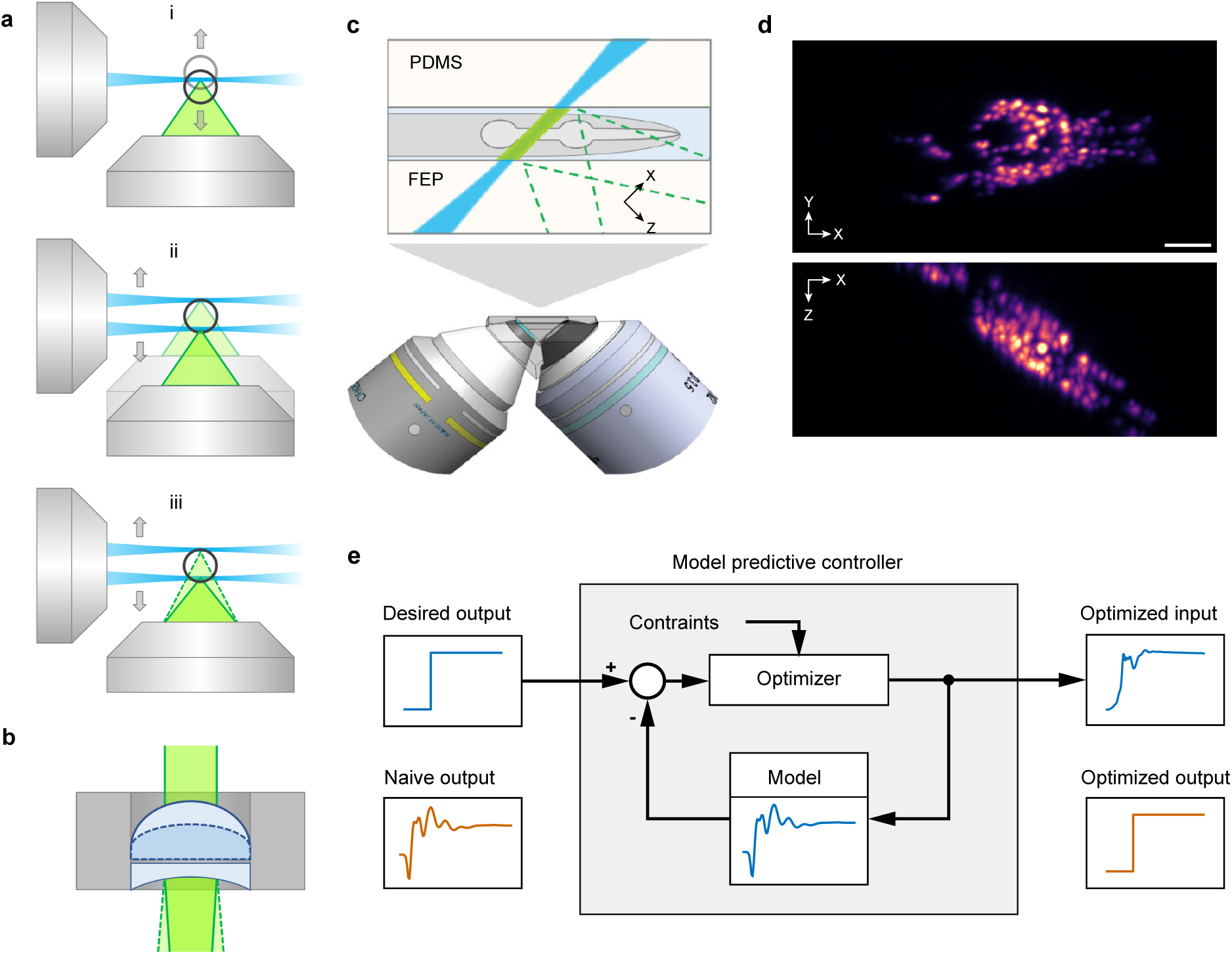
Open-top light-sheet microscope with electrically tunable lens (ETL) remote focusing under model predictive control. (a) Scan methods for volumetric light-sheet imaging. (i) In the simplest case, a sample is scanned through a static sheet. (ii) An actuated objective lens can follow a scanning light-sheet, which is faster than (i) but inertia limited and can cause problems with water immersion. (iii) Remote focusing optically shifts the focal plane using an active element like an ETL (b). This setup has less inertia and no moving parts at the sample. (c) Long-working distance open-top imaging is achieved using an asymmetric pair of objective lenses coupled with a water immersion fitting. This enables unobstructed imaging across a water-matched barrier such as FEP (above). (d) Maximum intensity projections along the Z (top) and Y (bottom) axes of a worm expressing a pan-neuronal nuclear-localized fluorescent protein positioned in a microfluidic channel as shown in (c). Scale bar 20 *µ*m. (e) Fast actuation of an ETL induces high frequency oscillation which slows response time. Performance is improved by using model predictive control to optimize drive signals. The controller iteratively optimizes the input signal to the ETL by minimizing simulated output error while obeying system constraints.

The rest of the microscope is divided into independent modules based on existing designs which can be modified to meet experimental needs (Supplementary Fig. 1). The excitation arm of the microscope follows the iSPIM design [24]. Excitation light from a fiber-coupled laser source (Coherent OBIS) is steered with a micromirror scanner (ASI) and passes through a tube lens and the illumination objective to the sample. While a scanned virtual light-sheet provides better image quality, we obtained best speed for the specific sCMOS camera used (Flash 4.0v3, Hamamatsu) using a static light-sheet formed by the addition of a cylindrical lens. The excitation light is flashed in time with the camera exposure to minimize motion artifacts. Emitted fluorescence is collected by the detection objective lens and focused to an intermediate image by the tube lens. The image is then relayed twice, first through a remote focus module and then through a custom image splitter and onto the camera. The remote focus module is configured as described by Fahrbach et al. [14] with the addition of a Dove prism placed in the infinity space after the ETL, which allows rotational alignment of the image. The image is then masked and relayed through the image splitter, which is designed to allow simultaneous imaging of up to four fluorescence channels spaced along the fast readout axis of the camera. This facilitates high-speed multichannel imaging tailored to our application: multichannel recording from *C. elegans* neurons primarily while restricted to a microfluidic channel.

### 3.2 Model-predictive control of tunable lens dynamics

Generating an empirical model for MPC optimization first required a way to accurately measure focal shift at high speed. To do this, we took advantage of the tilted imaging plane of our light-sheet microscope configuration to image a thin-line fluorescent target created by turning the light-sheet to a vertical orientation and illuminating a sample of 20 percent fluorescein (Fig. 2a). At such high concentration, fluorescence emission is dominated by absorption, resulting in a thin layer of excited fluorescence at the coverslip in the shape of the light-sheet cross-section [25][26]. This line intersects with the imaging plane at a single point, which shows in-camera as a brightness peak in the center rows of pixels that shifts linearly as the focal plane moves (Fig. 2b,c). By restricting the camera ROI to the center 16 rows of pixels, we were able to record at 5,000 frames per second and synchronize the analog drive signal to the camera readout clock.

Using this approach, we measured the response of the remote focus system to an input chosen to maximize the information available for system identification. The ideal signal for this purpose is usually some form of white noise – random input that is spectrally uniform and has maximum signal entropy. In practice, we preferred to use a pseudorandom binary m-sequence rather than Gaussian random noise, as the finite amplitude made it easier to ensure the input was within device limits at all times. We used an m-sequence with 15-bit word length, which at 5 kHz sampling rate provided about 6.5 seconds of non-repeating input with a guaranteed zero mean. While this run length is much more than required for identifying dynamics expected to settle within 20 ms, it was useful to capture additional, much slower dynamics we had previously observed, which may be the result of current-dependent heating and cause a hysteresis-like effect if not accounted for.

To probe system response, we measured the location of the line target brightness peak while delivering the m-sequence drive signal at a small amplitude of *±*0.1 V (Fig. 2d) around multiple center voltages. This approach allowed us to test for nonlinearity in the response while maximizing the likelihood of a good linear fit to each m-sequence trial. We measured and controlled the remote focus module, including both the ETL and voltage-controlled current source (VCCS) driving it, as a single unit. Therefore, system inputs are described here as voltages delivered to the VCCS over its input range of 0-5 V, which was approximately mapped to the ETL input current range of 0-300 mA. Each trial also included a sequence of voltage steps used to calibrate the line target readout to an equivalent steady-state voltage (Veq). This allowed us to correct for any errors due to non-uniformity in the fluorescence and made the input-output relationship a unitless ratio that could be scaled to calibrated physical units later. Each trial was also followed by a sequence of repeating square pulses to allow quick estimation of model fit to a step input. The switching rate of the analog output device (PCI-6733, National Instruments) is orders of magnitude faster than the intrinsic bandlimit of the measured system, making the probe signal close to ideal white noise.

We then used the m-sequence response data to build a linear, time invariant (LTI) model of the combined remote focus system. In order to capture dynamics at multiple timescales, we first estimated a 500 sample (100 ms) impulse response in MATLAB and then used MATLAB’s Model Reducer to reduce this to a 16th order state space model. The generated unit step response and frequency response magnitude of this model are shown in Figure 2e. The identified model captures features expected from manufacturer product specifications, including a strong under-and overshoot in the step response that settles in 15 ms and multiple high frequency resonances. Repeating the system identification at multiple offsets did reveal a nonlinear dependence on drive magnitude causing the resonant frequencies to shift (Supplementary Fig. 5). This high frequency oscillation is likely a higher order vibrational mode in the physical membrane that forms the lens, so it may be that the wave speed changes as the membrane is stretched. While this means an LTI model is insufficient to predict the response to all inputs, the identified model was excellent at predicting high frequency behavior around the center voltages on which it was tuned, and the model parameters were easily re-tuned to accurately model a step toward any set point. For the purpose of optimizing a single scan waveform, we found it sufficient to over-fit the model to the desired scan amplitude, but extension to a more general use could possibly be achieved using a collection of linear models or a linear parameter varying model to capture the frequency shift.

Finally, we used the generated model to construct a model predictive controller using the MATLAB Model Predictive Control toolkit. The controller includes the relevant limits of the system as constraints. Most importantly, the drive signal is hard limited to the input range of the current source, in this case 0-5 V, which forces the optimizer to produce solutions which are physically achievable. On the output side, we limit the minimum predicted output to 4 percent above the steady state zero per manufacturer recommendations to avoid damage to the ETL that can occur if the internal voice coil impacts the housing. The prediction horizon of the controller was set to cover 15 ms, which allows the optimizer to account for the full high-frequency response in its control solution. The controller is also configured to look ahead at the future reference signal, since the desired scan waveform is known before operation. This improves performance by allowing the controller to anticipate large maneuvers, such as sawtooth flyback, and begin making adjustments to minimize future error.

Using the MPC-optimized drive signal, we are able to eliminate undershoot and reduce the overshoot of the system step response from over 50 to less than 10 percent (Fig. 2f). Some high-frequency ringing appears to be unavoidable with a linear model; however, the 5 percent settling time was reduced to 5 ms (7 ms for 2 percent). In practical terms, this allows our light-sheet microscope to operate at the maximum speed allowed by camera and beam scanner limits. This performance improvement is consistent with the results of Iwai et al. [20] using sparse optimization. However, our MPC approach is substantially faster. Sparse optimization of a 40 ms sequence of focus steps was reported to take 30 seconds to compute. In contrast, MPC generated an optimized scan pattern for the full 9.7 second probe sequence in under 3.1 seconds. While we used the MPC as an offline optimizer here, this means our method is capable of real-time control. Additionally, while sparse optimization is a global method that requires the full reference signal in advance, MPC only actually requires advance knowledge of the reference signal over the prediction horizon, in this case 15 ms.

**Figure 2:**
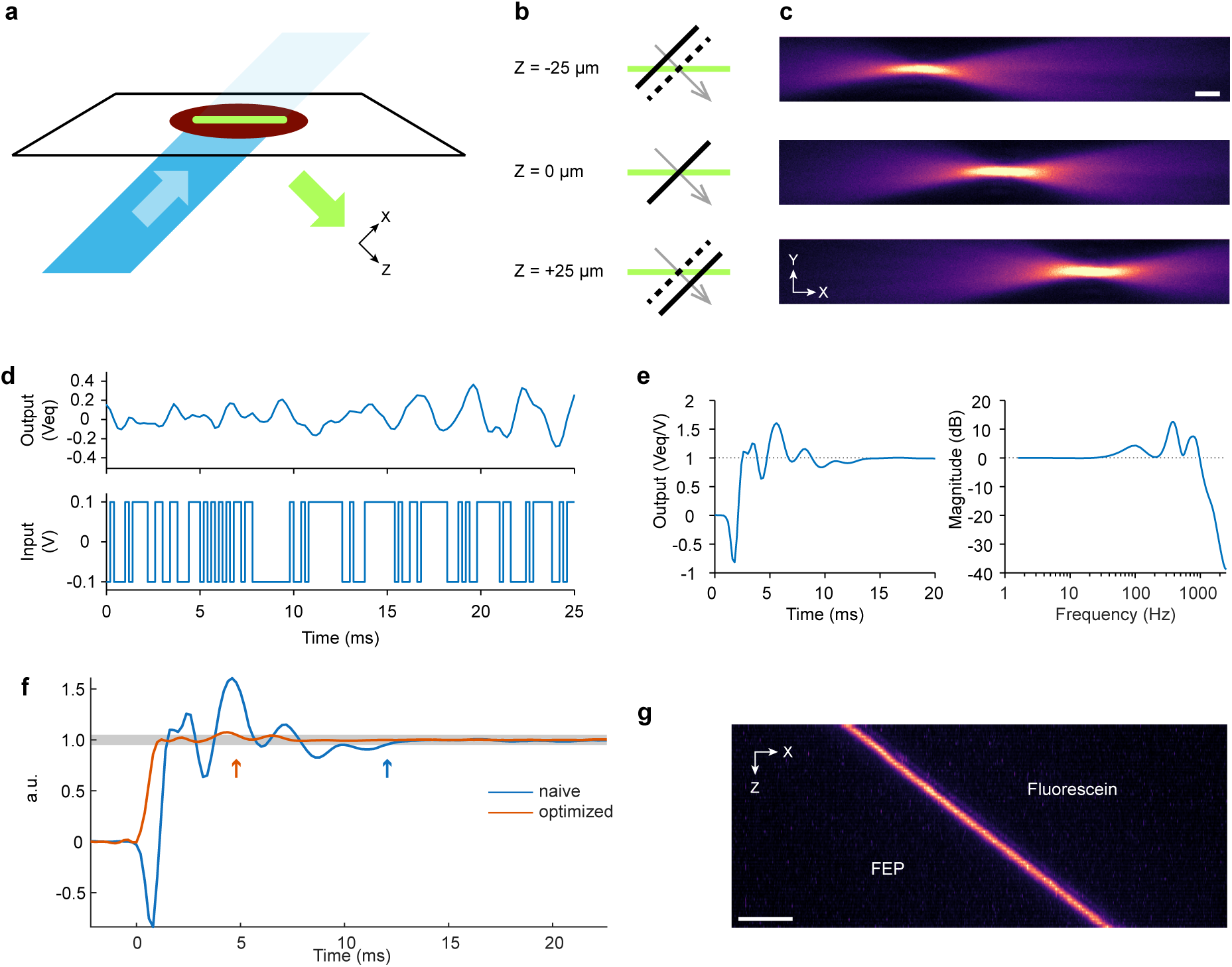
Measurement, modeling, and control of tunable lens dynamics. (a) Focal plane shift is measured using a thin-line fluorescent target, created by using a vertically oriented light-sheet to excite a layer of 20 percent fluorescein. (b) Focus shift changes the point at which this line intersects the focal plane, which is visible as a moving brightness peak in-camera (c). Scale bar is 10 *µ*m. (d) Measured focus position over time in response to a pseudorandom binary input at 5 kHz. Position is normalized to equivalent steady-state voltage (Veq). (e) Unit step response and frequency response of best fit linear time invariant model of remote focusing module dynamics. (f) Normalized measured responses to naive and MPC optimized step. The optimized drive signal avoids excessive movement and improves settling time. Traces are aligned to set t=0 to last time point within 5% of start point. Shaded area is *±*5% of target set point. Arrows indicate 5% settling time. Traces are averages of 10 repeated steps. Standard deviation from mean was ¡0.5%. (g) Y axis maximum intensity projection of fluorescence target in (a) using optimized volumetric imaging scan synchronized to light-sheet movement. Stack consists of 60 slices recorded at 10 vol/s. Scale bar 20 *µ*m

### 3.3 Fast whole-brain imaging of *C. elegans* enabled by tunable lens refocusing with model predictive control

As a first demonstration, we recorded the whole brain response of *C. elegans* under chemosensory stimulation delivered with microfluidics. The microfluidic device used (Fig. 3a) is a well-established design for chemical stimulation of the *C. elegans* head [22], which was bonded to an FEP substrate. The device was mounted by taping the FEP film to a simple 3D printed slide with a hole in the center, which was then placed in a standard slide mount insert on the motorized XY translation stage (Supplementary Fig. 4. We used a matching 3D printed alignment jig to ensure the microfluidic channel containing the worm was aligned with the imaging axis in order to prevent the PDMS channel walls from interfering with light-sheet formation or image collection. The configuration of objective lenses and immersion fitting provided sufficient clearance for free lateral movement and unobstructed access to the microfluidic device from above. Using remote focusing with our model predictive ETL control, we achieved multichannel volumetric imaging at a camera-limited 10 volumes per second at sufficient resolution for single-cell segmentation and tracking. Each volume consisted of 60 slices spaced approximately 1.25 *µ*m apart with a sawtooth scan pattern, which provides even temporal sampling between volumes. While higher volume rates were possible with fewer slices, this spacing was well matched to the thinnest light-sheet covering the full field of view, and we found that larger spacings reduced the separability of densely packed neurons near the nerve ring.

The worm strain used expresses pan-neuronal (*rgef-1p*) nuclear GCaMP6s and mNeptune and a sparse landmark label (*unc-47p* + *gcy-32p*) of nuclear CyOFP to aid in cell tracking and identification. These were imaged simultaneously in three channels (green, red, and deep red) arranged along the fast axis of the sCMOS camera (Fig. 3b). Simultaneous imaging of multiple fluorescence channels in this way is essential to maximize speed, but not without its own tradeoffs. The simple custom image splitter used lacks correction for chromatic defocus, making it impossible for all channels to be in sharp focus simultaneously. However, we found the amount of blurring acceptable for nuclear-localized labels with focus balanced between channels. Fluorescent proteins in general also have long-tailed emission spectra, so cross-contamination of fluorescence channels is expected. In this configuration, mNeptune emission was equally detected in the red and deep-red channels, while CyOFP was mostly confined to the red channel. We used the deep red for functional reference as it has the least included green fluorescence, and used the ratio of red to deep red fluorescence to identify landmark neurons.

The responses of *C. elegans* chemosensory to a wide array of odorants have been studied extensively, although usually as individual cells. Recent work using calcium imaging combined with microfluidics to characterize the sensory neuron responses to a wide array of chemosensory cues revealed that even single odorants are detected by an array of sensory neurons that encode the information as an ensemble [27][28][29]. We continuously imaged at 10 volumes per second while exposing a worm to five bouts of 10 seconds of benzaldehyde stimulus followed by 30 seconds of buffer. We used a 10*^−^*^4^ concentration of benzaldehyde as the stimulus, as it has been shown to evoke a reliable response that is compactly represented in the amphid sensory neurons [27]. This allows identification of the directly responding neurons by the combination of their response and relative location. Over 100 visible neurons were labeled and tracked in ZephIR [30]. Thanks to the light efficiency inherent to light-sheet microscopy, overall photobleaching was low, with mNeptune labels retaining nearly 70 percent of their initial brightness after 4.5 minutes of recording (Supplementary Fig. 6).

Calcium traces are reported as the ratio of green to deep red fluorescence, normalized by the mean of the dimmest 10 percent of time points to emphasize inhibition of tonically active neurons (Fig. 3c). As expected, we observed both stimulus-evoked activity and spontaneous activity, which was broadly distributed throughout detected neurons. Cross-correlation and hierarchical clustering of calcium traces revealed several distinct clusters of correlated and anti-correlated activity (Fig. 3d). We focus on three clusters with noteworthy patterns of activity, shown anatomically in Figure 3e. The first cluster (purple) demonstrates activity directly correlated with the stimulus, including the sensory neurons directly excited by benzaldehyde odor and closely connected interneurons. The other two larger larger clusters (orange and yellow) correspond to synchronized, spontaneous activity and are tightly anti-correlated with each other. This oscillating spontaneous activity is a common feature of *C. elegans* whole brain activity [31][32], and is expected to be dominated by interneurons and motor neurons that form mutually antagonistic circuits driving forward and backward crawling behaviors. Other clusters of spontaneous activity tend to join this pattern rather than act on their own timing, which may underlie hierarchical organization of behavior [32].

On the side of the worm closest to the detection objective (right), we identified three sensory neurons, AWA, AWC, and ASH, predicted to respond to benzaldehyde onset under these conditions [27] by their calcium activity and relative anatomical position. AWA and ASH were reliably excited by stimulus onset while AWC was reliably inhibited (Fig. 3f). As reported by Lin et al., we did not detect a response in AWC to stimulus removal after 10 seconds. Behaviorally, AWA and AWC are known as primary drivers of chemotaxis toward attractive odors. ASH, on the other hand, is a multimodal nociceptor which drives avoidance of noxious cues. All three are broadly tuned to detect volatile odors, and their antagonistic balance, along with input from the other chemosensory amphid neurons, allows the worm brain to represent its complex sensory environment in context dependent ways. While benzaldehyde is also expected to excite AWB on stimulus removal, we were unable to detect a cell with matching activity, which may be due to inconsistent expression of pan-neuronal promoters in the amphid sensory neurons.

While the calcium activity of neurons show consistent rapid response to stimulus onset, the long decay times seen are likely more reflective of the slow deactivation kinetics of the GCaMP6s fluorescent reporter used[33]. This asymmetry, responding to increases in calcium concentration within tens of milliseconds but decaying over seconds, is useful for reliably detecting excited calcium activity at lower imaging rates, but obscures inhibition events and graded responses. Taking advantage of the high temporal precision and signal-to-noise ratio of this recording, we performed a simple deconvolution of the sensory neuron activity traces using an exponential decay kernel to estimate underlying calcium dynamics for each sensory neuron response (Fig. 3g). Based on this analysis, estimated activity peaks quickly for both AWA and ASH before reaching a plateau and gradual decay. AWC shows a rapid fall in activity consistent with active inhibition by the stimulus and gradually recovers to a baseline seconds after stimulus removal. Most noteworthy is that the stronger AWA response to the first stimulus exposure could be explained by a sharp spike in calcium activity that rapidly returns to levels matching the following responses. AWA has been shown to rapidly respond to some odor gradients [34][35] with calcium-mediated action potentials, although this has only been demonstrated with high speed 2D imaging of AWA under an isolated label. While this analysis is insufficient to differentiate a true spike from a potential GCaMP nonlinearity, it demonstrates that higher imaging speeds may be essential to understanding the time sensitivity of distributed *C. elegans* sensory response and how such timing codes might affect downstream activity.

### 3.4 Reduced apparent motion from high-speed volumetric imaging

In addition to finer sampling of functional signals, increased imaging speed also reduces apparent motion between volumes. As soft-bodied invertebrates, *C. elegans* can contort their bodies in ways that interfere with anatomical tracking. This is even true for the neurons in the head ganglia of fully restrained worms, as pharyngeal pumping and muscle contraction cause internal elastic deformation. Restricting fluorescence excitation to a brief flash for each frame prevents motion blur during these events, but large, nonrigid movements can cause ambiguity when tracking cells over time.

To examine the effect of increased imaging speed on apparent cell movement, we recorded 20 vol/s whole brain fluorescence from a small day 1 adult worm constrained within a microfluidic channel but not fully immobilized (Fig. 4). Each volume consisted of 30 slices with 2.5 *µ*m separation. We tracked several reliably identified landmark cells distributed throughout the head using ZephIR (Fig. 4c). To simulate imaging at lower speed, we divided these results into six virtual 3.3 vol/s datasets by subsampling. The 3.3 volume rate is comparable to (or slightly faster than) the best imaging speed we have obtained in previous studies using a confocal microscope [36]. Unsurprisingly, the full-speed dataset shows smaller displacements between frames than subsampled versions. At full volume rate, 99.3 percent of displacements were less than 2 microns, the average radius of recorded nuclei, and 81.7 percent were less than the width of a single pixel (Fig. 4e). For subsampled recordings, this decreased to 87.4 and 32.7 percent, respectively.

**Figure 3:**
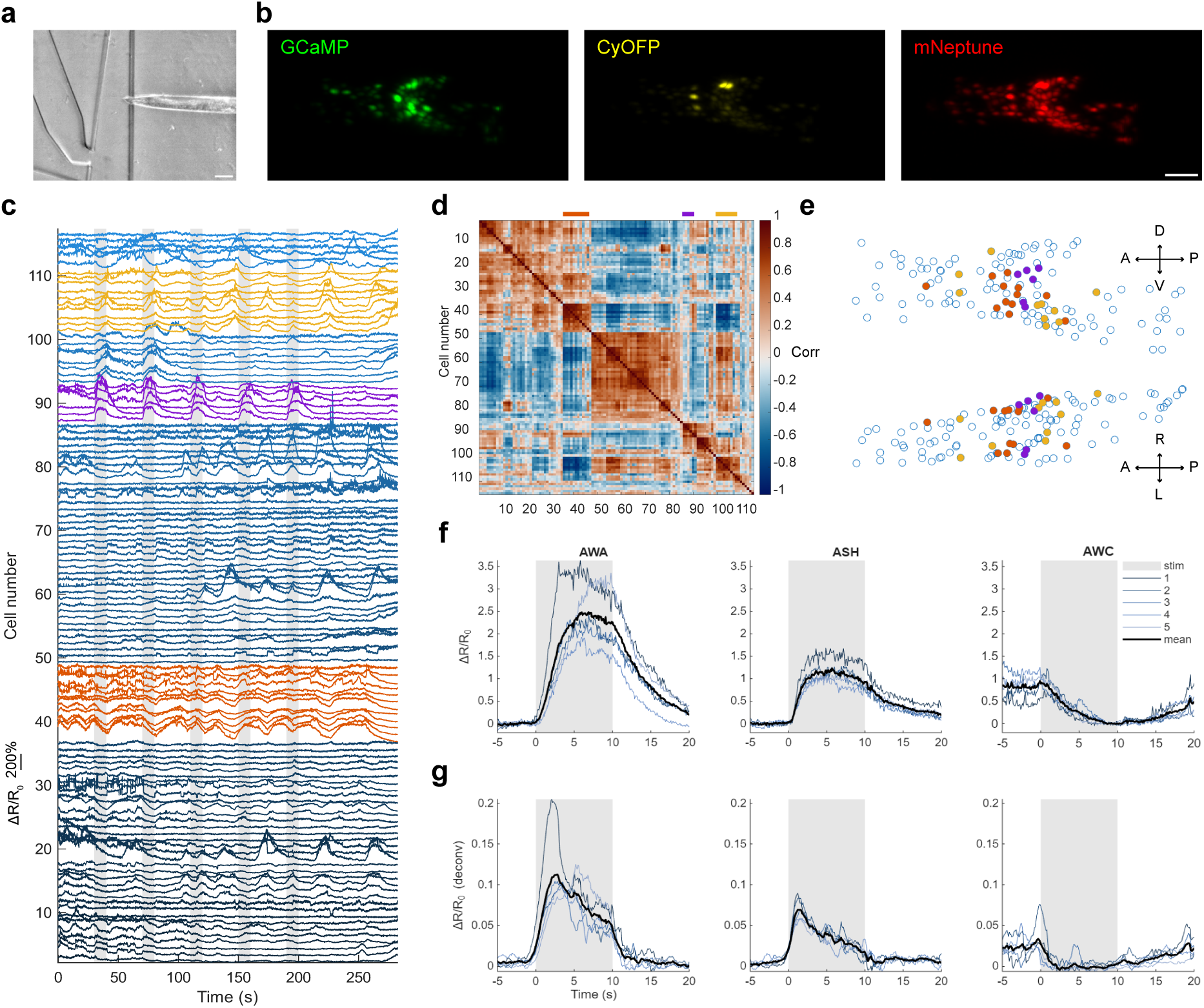
High speed imaging of whole brain response to benzaldehyde stimulus. (a) Infrared brightfield image of a worm in a chemical stimulation microfluidic device. Scale bar 100 *µ*m. (b) Rotated maximum intensity projections of green, red, and deep red fluorescence volumes acquired simultaneously. Scale bar 20 *µ*m. (c) Ratiometric calcium activity from tracked neurons, acquired at 10 volumes per second. Highlighted traces indicate stimulus-evoked (purple) and persistent, anti-correlated clusters of spontaneous activity (yellow and orange) identified by correlation analysis. Gray bars indicate stimulus periods. (d) Cross-correlation between activity traces sorted by hierarchical clustering. Heatmap shows normalized correlation coefficients of peak correlations. (e) Anatomical locations of highlighted neurons at same time point as (b). (f) Activity of identified sensory neurons on right side of worm, closer to the objective lens. Values for each trace are normalized to period before stimulus onset for AWA, ASH and period before stimulus removal for AWC. (g) Deconvolved calcium activity from (f) assuming exponential decay of GCaMP with 4 s time constant estimated from data.

Increasing the sampling rate also increases the rigidity of motion between frames. To show this, we simulated a simple translation-only registration step by subtracting the mean position of all tracked points within a volume before comparing positions between time points (Fig. 4d). After removing this bulk movement, displacements were 99.6 percent below 2 microns and 91.9 percent below pixel width for the full 20 vol/s dataset, but only 96.0 percent and 62.1 percent for 3.3 vol/s (Fig. 4f). In practice, we would expect this difference to be even larger for real recordings at 3.3 vol/s or slower, as the slices making up the volume would be sampled evenly over the full volume period. This separation in acquisition time between slices can cause even rigid motion to appear as non-rigid motion artifact in a recorded volume. These low displacement values, particularly after simple frame alignment, mean that tracking these cells between time points is unambiguous. This improves tracking results overall, and reduces the need for more computationally expensive methods.

### 3.5 Visualization of compartmentalized harsh touch response in PVD dendrites

Finally, we used our light-sheet microscope to observe the initiation of the harsh touch response in the *C. elegans* PVD neuron. The PVD neurons are unusual in the *C. elegans* nervous system for their extensively branched dendrite structure covering most of the body wall [37]. High intensity mechanical stimuli, referred to as harsh touch, elicit a cell-wide calcium response that triggers an escape response through synaptic signaling of downstream interneurons [38][39]. However, PVD also appears to be involved in maintaining body posture through non-synaptic, localized signaling and compartmentalized calcium activation has been observed in moving worms [40]. How these signals are integrated or isolated within the PVD neuron remains poorly understood, in part because of the difficulty posed by imaging calcium activity of this large, three-dimensional structure during any deformation required to probe its function. While the PVD dendrites are extremely thin (35-60 nm diameter in the smallest branches), the sparse structure means that image resolution can be sacrificed for speed as long as collection efficiency is sufficient.

Using our light-sheet microscope, we imaged calcium activity of PVD dendrites at 20 volumes per second during sustained compression in a microfluidic device used to probe harsh touch response [41] (Fig. 5a). Prior studies with this experimental preparation have generally focused on somatic response and required blanking the stimulation period due to the motion artifacts and loss of focus induced by deforming the worm [41][23][40]. We instead used a prolonged 5 second mechanical stimulus at 45 PSI and focused on a section of dendrites directly under the microfluidic valve providing the stimulus. We were able to continuously resolve several individual dendrite branches that remained in the field of view during mechanical stimulation, allowing us to observe the evolution of the induced calcium signal (Fig. 5b). The worm strain used expresses membrane localized variants of GCaMP7f and mScarlet in the PVD neurons (*ser2prom3* [40]), which were recorded simultaneously. Simultaneous imaging of a reference label like mScarlet was essential for segmentation and tracking, as baseline GCaMP fluorescence easily fell below the noise floor due to the usually weak expression of the fluorescent proteins under a PVD-specific promoter and small size of the dendrites. Ratiometric readout is also important for imaging during mechanical perturbation, as it should help to account for changes in local fluorophore density as dendrites are stretched.

**Figure 4:**
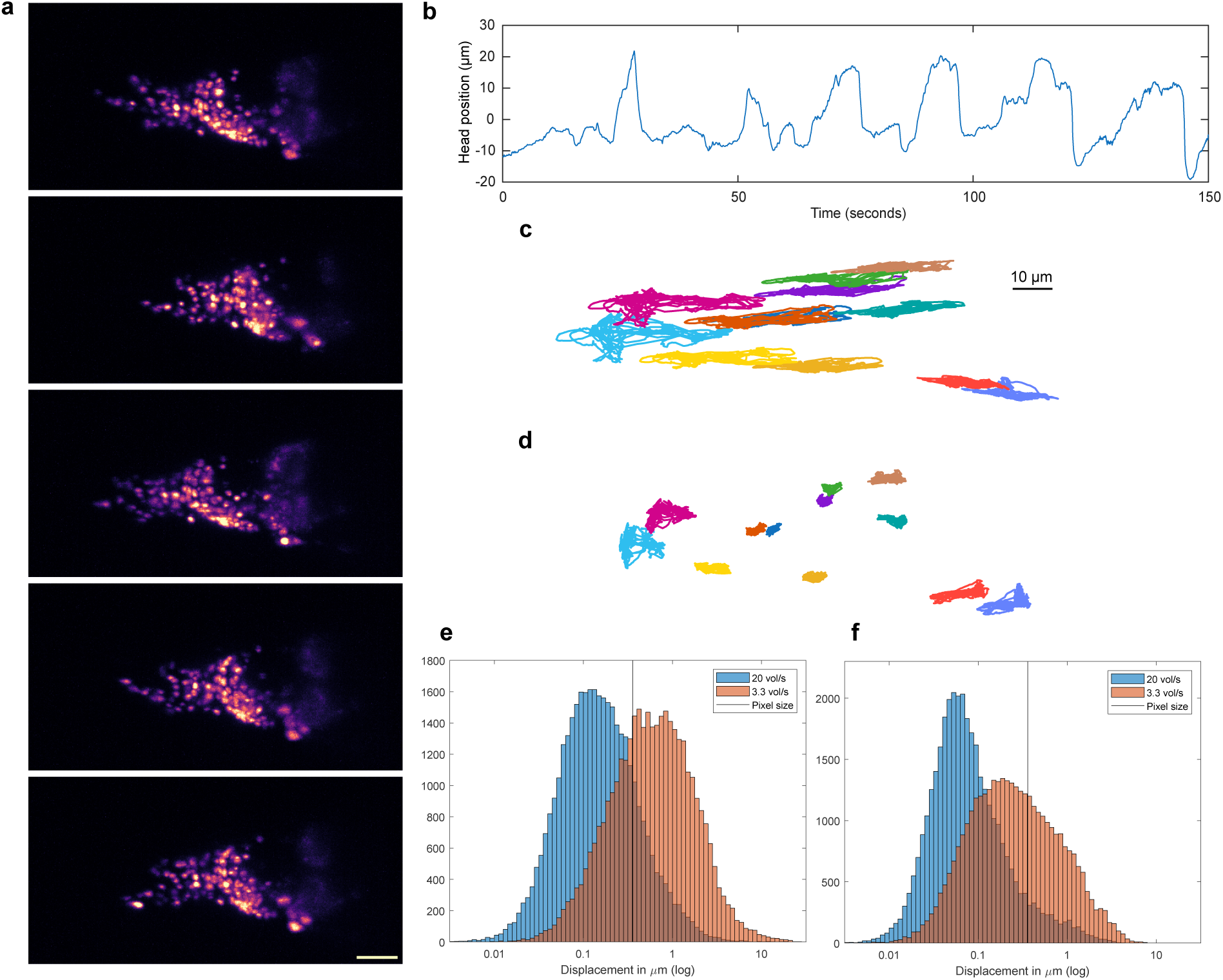
Apparent motion reduction with high speed volumetric imaging. (a) Example postures of an undersized worm in a microfluidic channel during whole brain imaging. Movement of the head causes elastic deformation of the head ganglia. (b) Bulk movement of the head in the channel over several minutes. (c) Paths traced by selected neurons over the recorded time period. (d) The same neuron traces after image alignment by simple translation for each time point. Residual movement highlights elastic deformation of the head. (e) Histogram of tracked cell displacements between time points. Lower speed (3.3 vol/s) within typical range of Piezo actuator scanning simulated by subsampling full speed (20 vol/s) data. Faster imaging reduced most cell movement to sub-pixel changes. (e) Adding a simple rigid registration step achieves subpixel alignment for more than 90 percent of cell displacements at 20 vol/s, but is less effective at lower speed due to non-rigid head movement.

As expected, the application of a harsh touch stimulus evoked calcium activity throughout the visible dendritic arbor (Fig. 5b). To investigate the spatial localization of this activity, we segmented the dendrites into smaller compartments by tracking points manually spaced along each branch in ZephIR and then tracing the brightest paths between points with Dijkstra’s algorithm. Extraction of calcium activity from these individual segments revealed considerable spatial variability in harsh touch response (Fig. 5c). Dendritic calcium activity appears to be bimodal, rapidly ascending to a plateau following an initial delay rather than gradually increasing throughout the stimulus interval. While this could be due to saturation of the GCaMP reporter, the spatial variation in the magnitude of these plateaus is not reflected in mScarlet fluorescence, suggesting a true difference in underlying calcium concentration. In the tertiary dendrites, this activation occurs earlier and reaches a higher magnitude at the distal ends, propagating as a wave toward the junction with the attached secondary dendrite. Calcium activity in the primary dendrite also varies spatially. Peak fluorescence and onset times vary non-monotonically along the dendrite, and may be compartmentalized between junctions with secondary branches. The secondary dendrites appear to have later and weaker calcium activity than the primary or tertiary dendrites on either side. These connective branches may serve as an active bottleneck that prevents subthreshold activation of tertiary dendrites from propagating and activating the soma, or they may simply restrict flow of calcium ions between compartments while membrane potential carries the noxious touch response. The mechanosensitive ion channels known to be involved in harsh touch sensation are expected to be highly selective for sodium ions, making the calcium activity seen likely due to secondary activation of voltage-gated calcium channels.

## 4 Discussion

In this work, we developed an open-top light-sheet with fast tunable lens remote focusing that improves scanning performance by optimizing drive signals through model predictive control. Using high-speed image-based measurement of focal plane movement with a spectrally rich probe signal, we generated a black-box LTI model which captured most of the ETL system dynamics. As the only modification made to ETL control was the choice of drive signal, this approach could be an in-place improvement to any application which uses tuneable lenses for high speed focusing. While ETL-based remote focusing may be inappropriate for applications demanding high resolution due to unavoidable aberrations, improved ETL control also has other applications than image focusing. For example, tunable lenses have also been used to provide fast axial positioning of focused excitation beams in axially swept light-sheet microscopy and multiphoton microscopy. These applications may benefit even more from model-based optimization, particularly axially swept light-sheets, which must reset at much higher frequency to synchronize with a rolling shutter.

**Figure 5:**
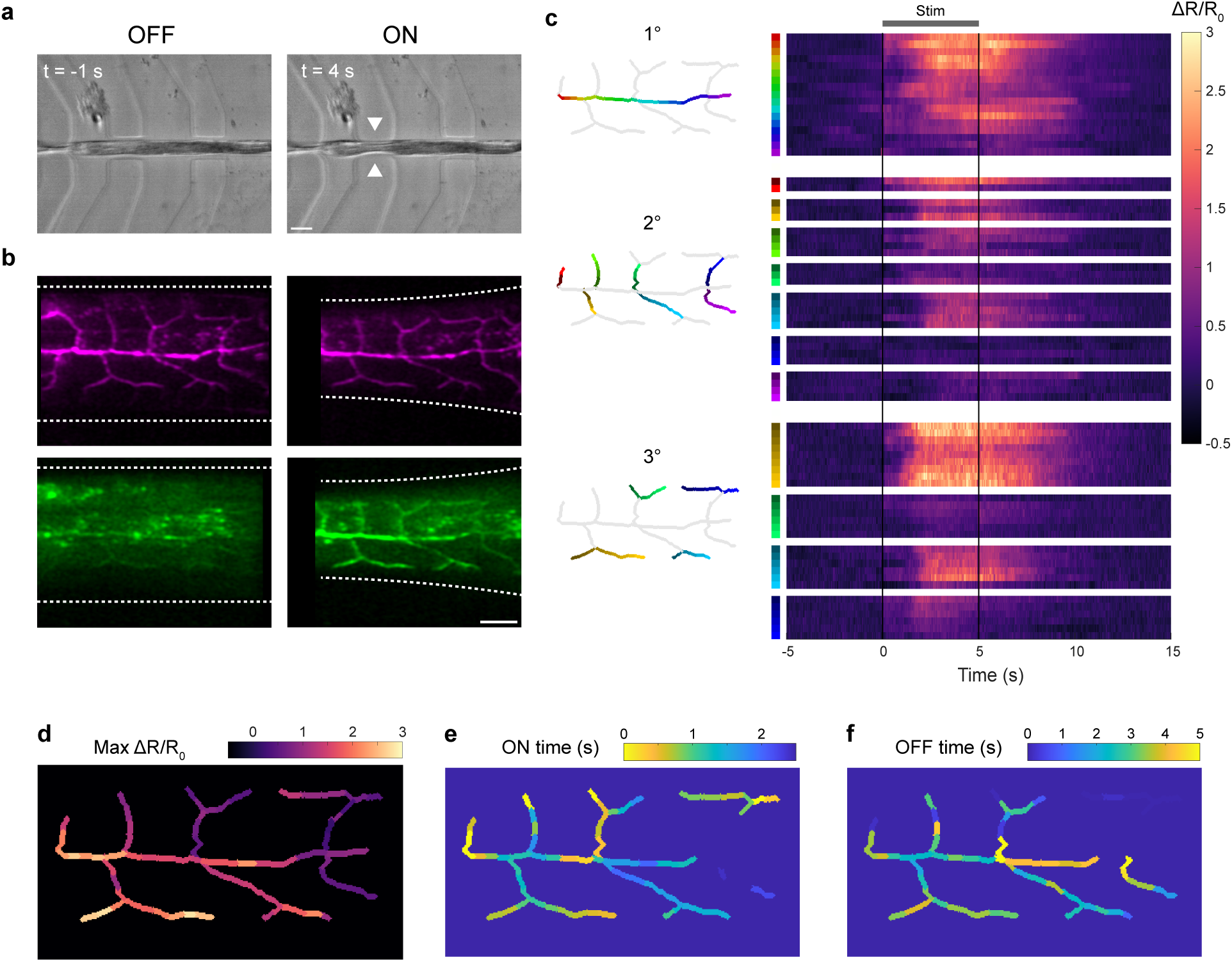
Compartmentalized activation of PVD dendrites. (a) Infrared brightfield images of a worm in a mechanical stimulation microfluidic device before (left) and during (right) anterior harsh touch stimulation. Scale bar 100 *µ*m (b) Depth-culled maximum intensity projections of mScarlet (top) and GCaMP7f (bottom) fluorescence at the same time points as (a). Scale bar 20 *µ*m (c) Calcium activity traces obtained from tracked segments of PVD dendrites, with positions shown in diagrams to the left. Harsh touch stimulation was applied from 0 to 5 seconds. (d) Spatial map of maximum fluorescence intensity for each segment. (e) Time after stimulus onset to reach half maximum intensity. (f) Time after stimulus removal to return below half maximum intensity.

Several limitations of this approach were acceptable for producing a consistent repeated scan pattern but may require additional work for applications requiring higher precision or fully generalized control. First, the lack of live feedback from the ETL means precise focal position is expected to drift, particularly due to the known temperature sensitivity of the device. For minutes-long recordings like those reported here, it is sufficient to compensate by adjusting light-sheet position at the start of imaging. Applications requiring stable positioning over longer timescales would need to at least apply temperature compensation. An online model predictive controller could also incorporate feedback of current ETL optical power, for example using an LED and photodetector. Also, our approach of overfitting the model for a specific waveform is only appropriate if high frequency excitation occurs at the same drive current. Fully generalized control, or even repeating waveforms with fast maneuvers at multiple positions, would need to incorporate the nonlinear dependence of resonant frequencies on absolute magnitude or accept slightly longer settling times. Nonetheless, MPC proved successful in generating optimized scan waveforms with performance comparable to far more computationally intensive methods [20] offline and with an LTI model, and it did so efficiently enough to be capable of real-time online control.

Remote focus not only improves speed compared to a moving objective, it simplifies the requirements of an open-top sample interface by eliminating the need to accommodate fast-moving parts. By combining remote focusing, an asymmetric pair of long working distance objective lenses, and a simple 3D printed water immersion fitting, we achieved a high-NA dual objective light-sheet microscope that can image into closed samples, including soft lithography microfluidics, provided the barrier is a water-matched material such as FEP. The choice of single-side acquisition with asymmetric objective lenses is primarily limited by the geometry of the lenses themselves. Given a symmetric pair of lenses with low enough profile, this sample interface and remote focus strategy could be adapted to a high-speed dual view design, which could achieve isotropic resolution through joint deconvolution [42].

Several limitations of the presented datasets are the result of camera-limited tradeoffs to maximize acquisition rate. Both the 10 and 20 volume per second scan configurations recorded 600 frames per second, with additional camera time needed for global exposure of pixels, additional discarded frames during sawtooth scan turnaround, and internal triggering delays. To maintain this rate, we under-sampled the image by adding a 0.5x magnification step at the image splitter, restricted the field of view to 256 rows of the camera, and imaged all fluorescence channels simultaneously across the camera fast axis. We also sacrificed some resolution by using a static light-sheet formed by cylindrical lens instead of a scanned light-sheet. While the camera used here supports virtual confocal recording of scanned light-sheet images using a rolling shutter, it would require halving the frame rate for the same field of view. Any of these image quality reductions could be traded back for reduced speed with this configuration, or both speed and resolution could be improved by using faster, more specialized, or multiple cameras for fluorescence detection. However, thanks to fast remote focusing and the independence of light-sheet formation and imaging in a dual objective system, these tradeoffs can be reconfigured on a per-experiment basis to take maximum advantage of available hardware.

## 5 Methods

### 5.1 Light-sheet microscope setup

The sample interface consists of an asymmetric orthogonal pair of water-immersion objectives: a 10x/0.4NA illumination objective (10X Nikon CFI Plan Fluorite) and 40x/0.8NA detection objective (40X Nikon CFI APO NIR). The objective pair is mounted to a 150 mm radius goniometer stage (Edmund Optics) and tilted to maximize working distance. Water immersion is maintained by a small fitting connected to a syringe with polyethylene tubing. The fitting was designed in SOLIDWORKS using freely available part models of the objectives lenses (obtained from Thorlabs) and 3D printed in Siraya Tech tenacious clear resin with an Original Prusa SL1S printer. Above the objective pair, samples are mounted as in a conventional inverted microscope on a motorized XY stage (MS2000, ASI) supported by a synchronized pair of motorized Z stages (Dual-LS-50-FTP, ASI).

Combined output from multiple lasers (OBIS 488nm LX 100mW, OBIS 561nm LS 80mW, OBIS Galaxy beam combiner, Coherent) is fiber coupled to a micro-mirror scanner (MM-SCAN-1.2, ASI). An adjustable static light-sheet is formed by addition of a free space half-cylindrical telescope between the collimator and scanner, consisting of a 50 mm cylindrical lens (ACY254-050-A, Thorlabs) in a rotating mount, an iris at the focal point for adjusting light width, and 100 mm spherical lens (AC254-100-A, Thorlabs). Light-sheet NA is adjusted with an iris in the micro-mirror scanner. The output scanned beam is directed through a 400 mm tube lens (AC254-400-A, Thorlabs) to the 10x/0.4NA illumination objective (Supplementary Fig. 1).

Emitted fluorescence is imaged through the detection objective, an excitation blocking filter (ZET488/561nm, Chroma), and 200 mm tube lens (AC254-200-A, Thorlabs) to an intermediate image plane, which is relayed through the remote focus module and image splitter to the camera. The remote focus module is a 4f relay formed by two 150 mm lenses (AC254-150-A, Thorlabs). An electrically tunable lens (EL-10-30-TC-VIS-12D, Optotune) is placed at the center of this relay and conjugate to the back focal plane of the detection objective. The ETL is driven by a voltage-controlled current source (T-Cube LED driver, Thorlabs). A -75 mm offset lens (ACN254-075-A, Thorlabs) is placed before the ETL to balance ETL-driven focus offset around the natural focal plane of the objective. A Dove prism (PS992M-A, Thorlabs) is positioned immediately after the ETL on a precision rotation mount to allow precise rotational alignment of the imaging field of view and the image splitter. The following intermediate image is masked and relayed again with a 200 mm lens (AC254-200-A, Thorlabs) into a series of two dichroic mirrors (FF560-FDi01 and FF640-FDi01, Semrock) and a turning mirror, which splits the image into green, red, and deep red paths. Each path includes an emission filter (FF03-525/50, FF01-608/54, and FF01-698/80 respectively, Semrock) and a small kinematic mirror to recombine the paths with a slight offset and allow precise positioning on the camera detector. Finally, the split images are focused onto the sCMOS camera (ORCA-Flash 4.0v3, Hamamatsu) with a 100 mm fixed focal length camera lens (86-410, Edmund Optics). The combination of 200 mm and 100 mm lenses de-magnify the final images 0.5X.

A separate near-infrared brightfield imaging path is positioned above the stage for animal tracking and microfluidic device monitoring. This consists of a long working distance 10x/0.28NA air objective (10x M Plan Apo, Mitutoyo), 830 nm longpass filter (FGL830, Thorlabs), 100 mm tube lens (AC254-100-A, Thorlabs), and CMOS camera (DCC3240N, Thorlabs). Brightfield illumination is provided by a 900 mW 850 nm LED (M850L3, Thorlabs), coupled to a gooseneck light guide positioned under the stage.

All hardware is controlled by a combination of an ASI Tiger controller (TG16 BASIC) and precision analog output device (PCI-6733, National Instruments). Image acquisition was coordinated using custom LabView code, with live volumetric image display provided by Napari by using a shared memory buffer. Before each acquisition, the scan parameters are used to generate a reference waveform which is passed to the model predictive controller in MATLAB to generate an optimized analog output.

### 5.2 Microfluidic device fabrication

Devices were fabricated in two layers using standard soft lithography methods. A soft bottom layer was made by spin coating 23:1 PDMS (Sylgard 184, Dow Corning) onto a feature master. A separate rigid backing layer was made by pouring 10:1 PDMS onto a blank silicon wafer. Both layers were baked at 70*^◦^*C for 30 minutes until rigid but sticky. The backing layer was then cut and placed on top of the feature layer and remaining space was filled with extra 10:1 PDMS. This was then cured overnight in a 70*^◦^*C oven.

After curing, devices were cut to size, and holes were added using biopsy punches (18 gauge for worm inlet and outlet, 19 gauge for valves). Strips of 50 or 125 micron thick Type-C Teflon FEP film were cut to size and ultrasonic cleaned in isopropyl alcohol for 10 minutes. Clean FEP strips were stored in deionized water for later use. PDMS devices were treated with a plasma wand for 30 seconds then placed firmly onto the hydrophilic side of a clean FEP strip. Devices were then thermally bonded by placing on a cold hot plate, heating to 200*^◦^*C for 30 minutes under pressure provided by a metal block, and removing once cool.

### 5.3 *C. elegans* strains and maintenance

*C. elegans* were cultured using standard maintenance methods on nematode growth medium (NGM) plates seeded with *E. coli* OP50 lawns [43]. Strain GT408 (whole-brain imaging, Fig. 3&4) was obtained by crossing ZM9624 (pan-neuronal GCaMP6s and mNeptune) with GT298, a landmark strain expressing CyOFP in 12 neurons, from previous work [36].

To record localized activation of the PVD neuron (Fig. 5), we generated strain GT367, which expresses membrane-targeted GCaMP7f and mScarletI under control of the *ser2prom3* promoter (GT367: *unc119(ed3) III; aEx46[ser-2p3b::GCaMP7F::ras-2CAAX::SL2::mScarlet-I::ras-2CAAX (50ng/ul) + pDSP2(Cbrunc-119(+) (50ng/ul)]*). To obtain GT367, *ser2prom3* was PCR amplified from genomic DNA and assembled to NotI digested linear pSiM4 backbone [44] using the NEBuilderHiFi DNA assembly master mix (New England Biolabs). The resulting plasmid, pSiM8 (*ser2prom3::GCaMP7f::ras-2CAAX::SL2::mScarlet-I::ras-2CAAX*) was injected into QL74 (*unc-119(ed3)*) at a concentration of 50 ng/*µ*L, in addition to 50 ng/*µ*L of pDSP2 (*unc-119* rescue).

The following promoters were used to amplify *ser2prom3* from genomic DNA: Forward 5’-CAGTCCAGT TACGCTGGAGTCTGAGGCTCGTCCTGAATGATATGCcgaaacgctgtcgactt-3’ and Reverse 5’-TCATA GAGGCCATTCCGTGGTGGTGGTGGTGGTGGGATCCCATtatgtgttgtgatgtcacaaaaatatgccaaaatg-3’.

All worms imaged were Day 1 or 2 adults obtained by either picking L4 larvae 24-48 hours before imaging or using a timed hatch-off procedure. For the timed hatch-off, adults and larvae were rinsed off from a mixed population plate, the remaining eggs were allowed to hatch for 3-4 hours, and L1 larvae were collected and grown at 20*^◦^*C for 3-4 days (Day 1 - 2 of adulthood) prior to imaging.

### 5.4 *C. elegans* functional imaging experiments

Functional imaging experiments were performed as previously described using a custom pressure control apparatus and MATLAB code to control mechanical and/or chemical stimulus delivery [22] [23]. For chemical stimulus experiments, the stimulus was prepared by diluting benzaldehyde to a concentration of 10 *^−^*^4^ (v/v) in NGM buffer (3 g/L NaCl, 1 mM CaCl_2_, 1 mM MgSO_4_, 25 mM KPO_4_) and adding 2 *µ*M fluorescein. All other imaging device inlets contained NGM buffer.

### 5.5 Image processing

All image volumes were denoised with a Butterworth low-pass filter and 2D aligned with the red channel before tracking. Whole brain recordings were annotated, tracked, and extracted in ZephIR[30] in raw orientation. Calcium traces are reported as the ratio of green to deep red fluorescence, normalized by the mean of the dimmest 10 percent of time points to emphasize inhibition of tonically active neurons. Regularized deconvolution of select traces was performed with the built-in deconv function in MATLAB 2025a.

PVD tracking was also performed in ZephIR, but with 2D images that were rotated and depth-culled to minimize the influence of autofluorescence. First, each volume was rescaled to isotropic voxel size and affine transformed to align the XY plane to the primary dendrite. Target Z depth was identified by the first slice to reach half maximum average intensity, and a 10 *µ*m span was maximum intensity projected. These projections were annotated and tracked in ZephIR by placing point labels along dendrites and at all branch points. After tracking, paths between adjacent points in each volume were traced in MATLAB using Dijkstra’s algorithm. Briefly, edges between all adjacent pixels were represented as an undirected graph, with edge weights between adjacent pixels were assigned based on fluorescence intensity. For an adjacent pair of pixels *x_i_*and *x_j_*:

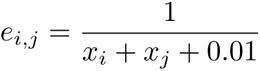

The resulting paths were converted to a segmentation mask by labeling the pixels corresponding to each point in the shortest path followed by grayscale dilation. Calcium traces were computed by taking green-t-red ratio of the mean of the brightest 15 pixels for each segment. Traces were debleached and normalized by taking log of the data, subtracting a linear fit to the first and last 100 samples, and exponentiating back.

## 6 Supplementary Figures

**Supplementary Figure 1:**
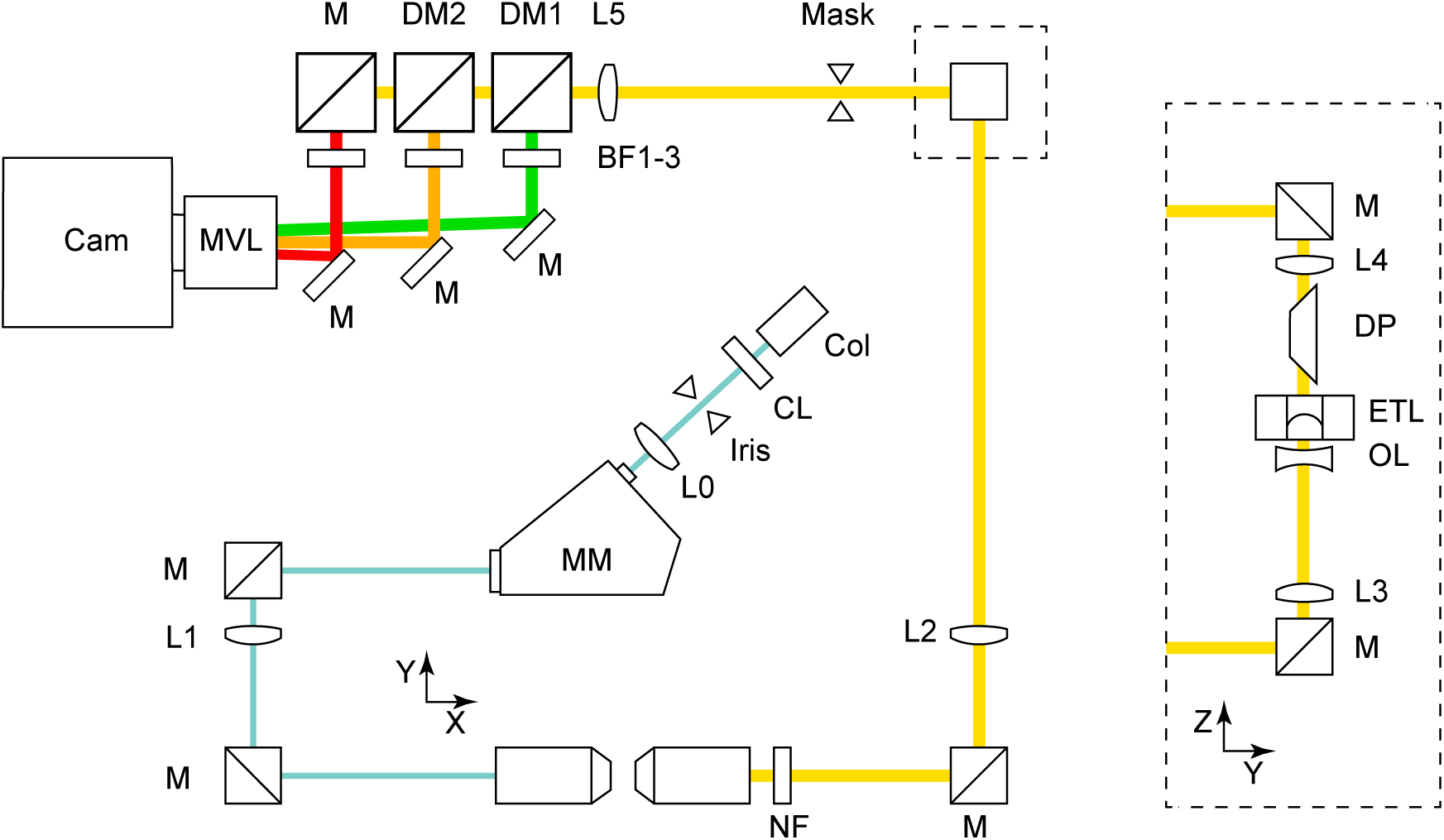
Light sheet microscope optics diagram (not to scale). Excitation path: Combined lasers leave collimator (Col) into optional static light sheet former formed by cylindrical (CL) and spherical (L0) lens pair. An iris at the cylindrical focal point selects light sheet width. Light sheet NA is selected with iris inside micromirror scanner (MM). Scanned beam passes through turning mirrors (M) and 400 mm tube lens (L1) to excitation objective. Detection path: Emitted fluorescence is captured by the detection objective, passes through a notch filter (NF) to block excitation light, and focused by a 200 mm tube lens (L2). The intermediate image is relayed through the remote focusing module (left inset) by two 150 mm relay lenses (L3-4). Remote focus is achieved by an electrically tunable lens (ETL) placed at the focal point between L3 and L4 and conjugate to the back focal plane of the objective. A -75 mm offset lens (OL) centers the optical power curve of the ETL. The image is rotationally aligned by a Dove prism (DM) to correct for tilt of objective. The module is oriented vertically to avoid gravity aberration in ETL. Image splitter: Second (static) intermediate image is masked and relayed again through lens L5. The image is split into spectral channels by a bank of dichroic mirrors DM1-2 and and bandpass filters (BF1-3). Channels are reflected to a machine vision lens (MVL) and individually aligned along fast axis of sCMOS camera (Cam) by small kinematic mirrors.

**Supplementary Figure 2:**
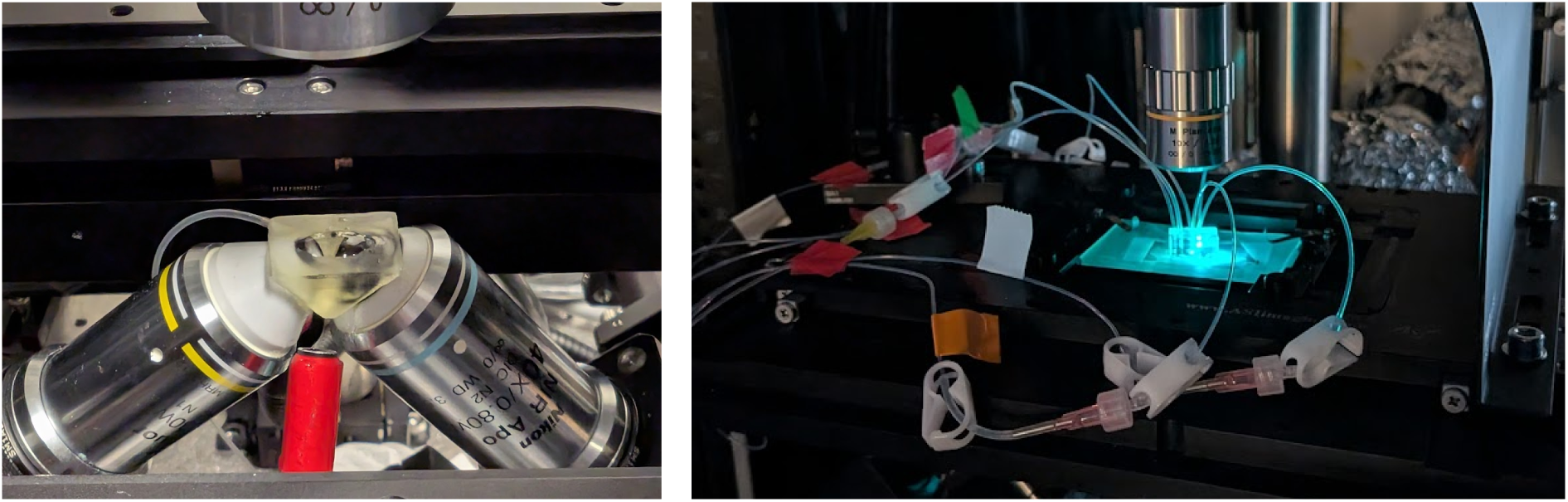
Open-top light sheet microscope sample interface. Left: objective lenses and 3D printed water immersion fitting, visible with stage insert removed. Fitting is filled with water by syringe via attached tubing. Right: Microfluidic device mounted on stage for imaging, illuminated by excitation laser. Open-top configuration allows free access for tubing. Objective lens for IR brightfield monitoring is visible above.

**Supplementary Figure 3:**
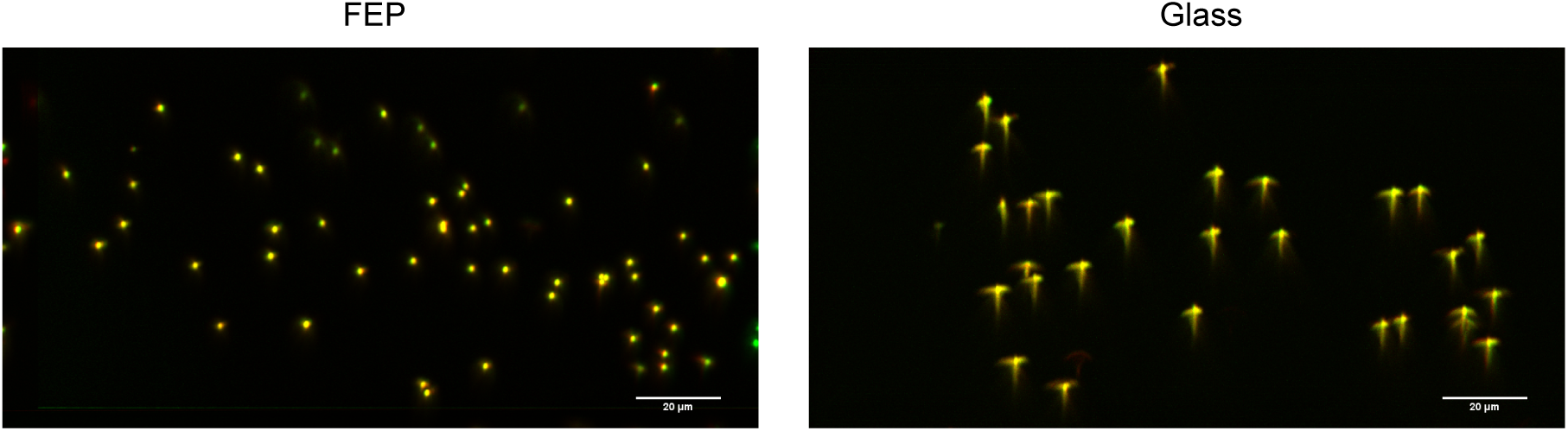
An index-matched material such as FEP is necessary for oblique angle imaging. Maximum intensity projections of 1 *µ*m fluorescent beads on FEP film (left) and a glass coverslip (right). Scale bars 20 *µ*m.

**Supplementary Figure 4:**
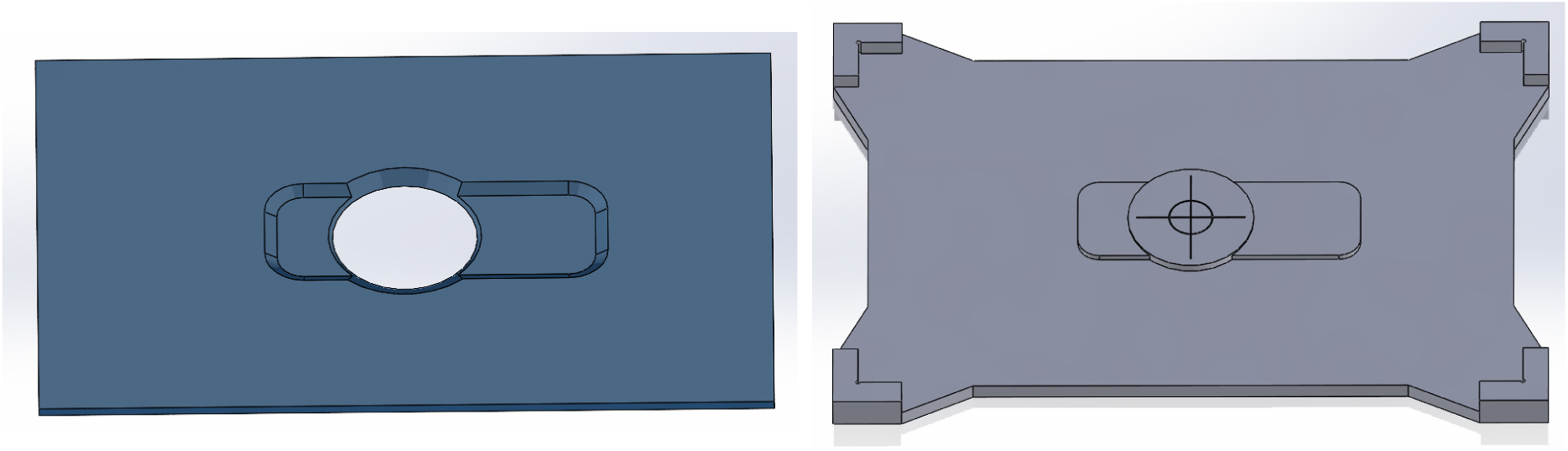
Renders of simple mount plate and alignment jig used to support FEP-based microfluidic device on XY stage.

**Supplementary Figure 5:**
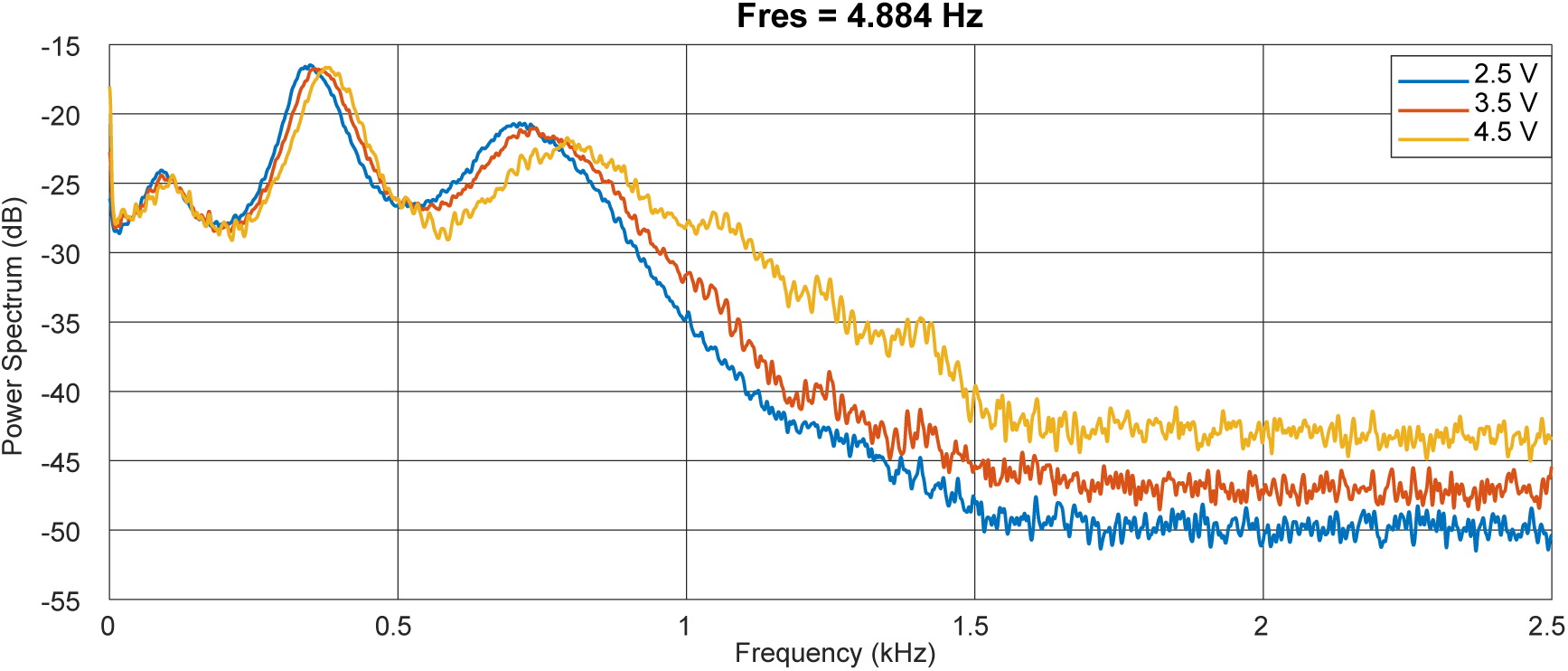
Resonant frequencies shift as center voltage increases. Power spectra computed from m-sequence recordings at center voltage *±*0.1 V.

**Supplementary Figure 6:**
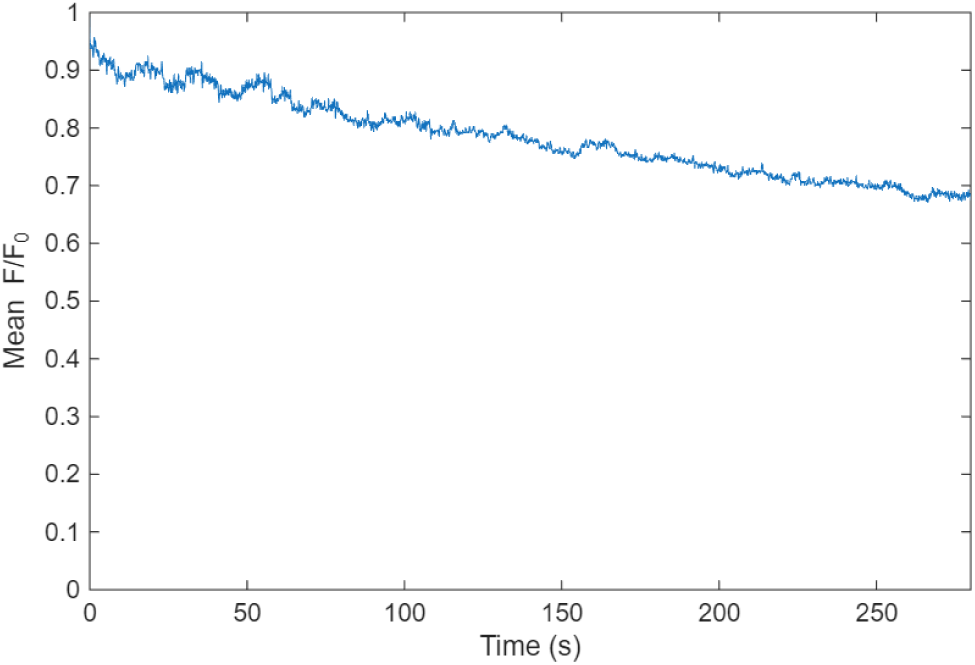
Photobleaching of mNeptune labels in recording corresponding to Fig 3 of the main text, normalized to the first time point. Recorded at 10 vol/s.

